# Substrate stiffness and cellular microenvironment regulate cell and junction mechanics in iPSC-derived brain microvascular endothelial cells

**DOI:** 10.64898/2026.01.06.698052

**Authors:** Li Yan, Udit Gupta, Haitao Wang, David S. Schrump, Kimberly M. Stroka

**Author notes:** Co-Corresponding authors Email addresses (K.M. Stroka); (L. Yan).

## Abstract

The blood–brain barrier (BBB) is a mechanically responsive interface that protects the central nervous system. Brain tissue exhibits region-specific stiffness that evolves throughout development and is altered in aging and various neurological diseases. These stiffness changes are increasingly recognized as key modulators of endothelial cell behavior and BBB integrity. However, the mechanisms by which brain endothelial cells sense and adapt to variations in their mechanical microenvironment remain poorly defined. Moreover, how mechanical cues interact with cellular signals from astrocytes and pericytes to modulate endothelial mechanics and junctional organization has been largely unexplored. Here, we demonstrate spatial regulation of subcellular mechanics in human iPSC-derived brain microvascular endothelial cells (iBMECs) in response to physiologically and pathologically relevant substrate stiffness (1–194 kPa). Using atomic force microscopy, we quantified Young’s modulus at three distinct cellular regions—tricellular junctions, bicellular junctions, and cell bodies. iBMECs cultured on compliant substrates (1, 2.5, and 15 kPa) exhibited pronounced mechanical polarization, characterized by significantly elevated stiffness at tricellular regions compared with bicellular regions and cell bodies. This spatial organization was lost on supraphysiological stiffness (194 kPa), which reduced overall cell stiffness and eliminated regional distinctions. Co-culture with astrocytes and pericytes decreased global stiffness but preserved the dominant reinforcement at tricellular regions. In contrast, exposure to metastatic breast cancer cells abolished junction polarization at tricellular regions and suppressed stiffness across all regions, particularly on soft substrates. These findings reveal that BBB endothelial mechanics are regulated by both matrix stiffness and BBB cell context in a region-specific manner. This work provides new insight into how physical and cellular cues shape BBB structure and function, with implications for understanding barrier disruption in neurological disease and metastasis.

## Introduction

The brain is one of the softest organs in the human body, and its development and homeostasis are highly influenced by mechanical cues, including extracellular matrix (ECM) stiffness^1^. Brain tissue exhibits considerable structural heterogeneity, with regional variations in stiffness that reflect both physiological and pathological states^1, 2^. Changes in brain stiffness are increasingly recognized as important biomarkers of disease progression. For instance, significant tissue softening has been observed in neurological disorders such as multiple sclerosis^3^ and Alzheimer’s disease^4^, while elevated stiffness is associated with low-grade gliomas and metastatic brain tumors^5^. The brain stiffness alterations are closely linked to the regulation and function of the blood-brain barrier (BBB). The BBB, as a mechanosensitive structure, can change its physical properties by sensing its surrounding microenvironment ^6, 7^.

The BBB is a highly selective and dynamic interface that preserves central nervous system homeostasis^8^. It is primarily composed of brain microvascular endothelial cells (BMECs), which are supported and regulated by astrocytes and pericytes^9^. BMECs express specialized junctional complexes that tightly regulate paracellular permeability^10^. Increasing evidence indicates that ECM stiffness modulates BBB permeability by influencing these junctional structures^7, 11^. Specifically, ECM stiffness regulates the assembly of junction proteins through mechanical tension ^12^. Our previous work has demonstrated that culturing primary human BMECs on soft substrates reduces cellular contractility, which correlates with enhanced tight junction coverage and barrier integrity ^13^. Additionally, we have also shown that increased substrate stiffness enhances leukocyte transmigration. In particular, stiffer matrices promote neutrophil transmigration across human umbilical vein endothelial cells stimulated with tumor necrosis factor-alpha^14^ or oxidized low-density lipoprotein^15^. Furthermore, our recent findings further show that substrate stiffness locally regulates BBB permeability by altering junctional organization, particularly at tricellular junctions^6^. These results suggest that substrate stiffness not only affects overall barrier integrity but also governs the spatial distribution of junctional proteins in a region-specific manner.

Junctional proteins are closely integrated with the cytoskeletal network, governing the mechanical architecture of endothelial cells^16^. Stiffness is a well-recognized regulator of endothelial cell mechanics, influencing cellular behavior under both physiological and pathological conditions^17^. When mechanosensitive proteins are subjected to external forces, cells often respond by increasing their stiffness to resist deformation^18^. For example, endothelial cells exert increased cell-matrix traction stress when residing on stiff hydrogels^19^. The spatial regulation of junctional organization by substrate stiffness, as shown in our previous findings^6^, further suggests that regional variations in stiffness may play a role in modulating endothelial structure and function. However, the mechanisms by which stiffness influences junctional organization and its implications for physiological and disease states remain incompletely understood. Furthermore, the mechanical phenotype of BMECs is shaped not only by ECM stiffness but also by interactions with neighboring cells, particularly astrocytes and pericytes. Despite their known role in barrier regulation, how astrocytes and pericytes influence the mechanical properties of BMECs in response to varying ECM stiffness remains largely unexplored. Importantly, the connection between cellular mechanical changes and barrier dysfunction in health and disease is only beginning to be elucidated.

In this study, we investigate how ECM stiffness and interactions with astrocytes and pericytes modulate the mechanical properties of human iPSC-derived BMECs (iBMECs). We focus specifically on region-specific stiffness at subcellular regions using atomic force microscopy (AFM) and tunable hydrogel substrates that mimic brain stiffness. We show that pericytes and astrocytes contribute to region-specific modulation of iBMEC mechanics, with pronounced effects at tricellular junction regions. Furthermore, we show that brain-seeking breast cancer cells (MDA-MB-231-BR) selectively disrupt stiffness at tricellular junction regions, particularly on soft matrices, thereby enhancing their incorporation into the iBMEC monolayer. These findings uncover a previously unrecognized, mechanically-mediated mechanism of BBB regulation and metastatic vulnerability, highlighting the importance of region-specific mechanobiology in both health and disease.

## Results

### Matrix stiffness and co-culture with astrocytes and pericytes modulate iBMEC morphology and mechanical properties

To study the subcellular mechanical landscape of iBMECs and how it is modulated by their mechanical and cellular microenvironment, we established an in vitro BBB model composed of iBMECs co-cultured with astrocytes and pericytes on polyacrylamide (PA) hydrogels of tunable stiffness (Figure 1A). Immunofluorescence of requisite markers confirmed the identity of pericytes, astrocytes, and iBMECs (Figure 1B). To assess regional differences in mechanical properties, we performed AFM on iBMECs cultured under defined stiffness conditions. AFM force mapping generated both topographical and Young’s modulus maps (Figure 1C). Topography images indicate changes in cell morphology and monolayer architecture. Young’s modulus maps show regional variation in stiffness. To quantify these regional differences, we implemented a computational segmentation approach using a custom MATLAB pipeline. This method allowed us to define and extract stiffness values from specific regions within the iBMECs: tricellular junctions, bicellular junctions, and the cell body (Figure 1D). Using this approach, we identified the patterns of regional mechanical heterogeneity of iBMECs across different stiffness conditions.

**Figure 1.**
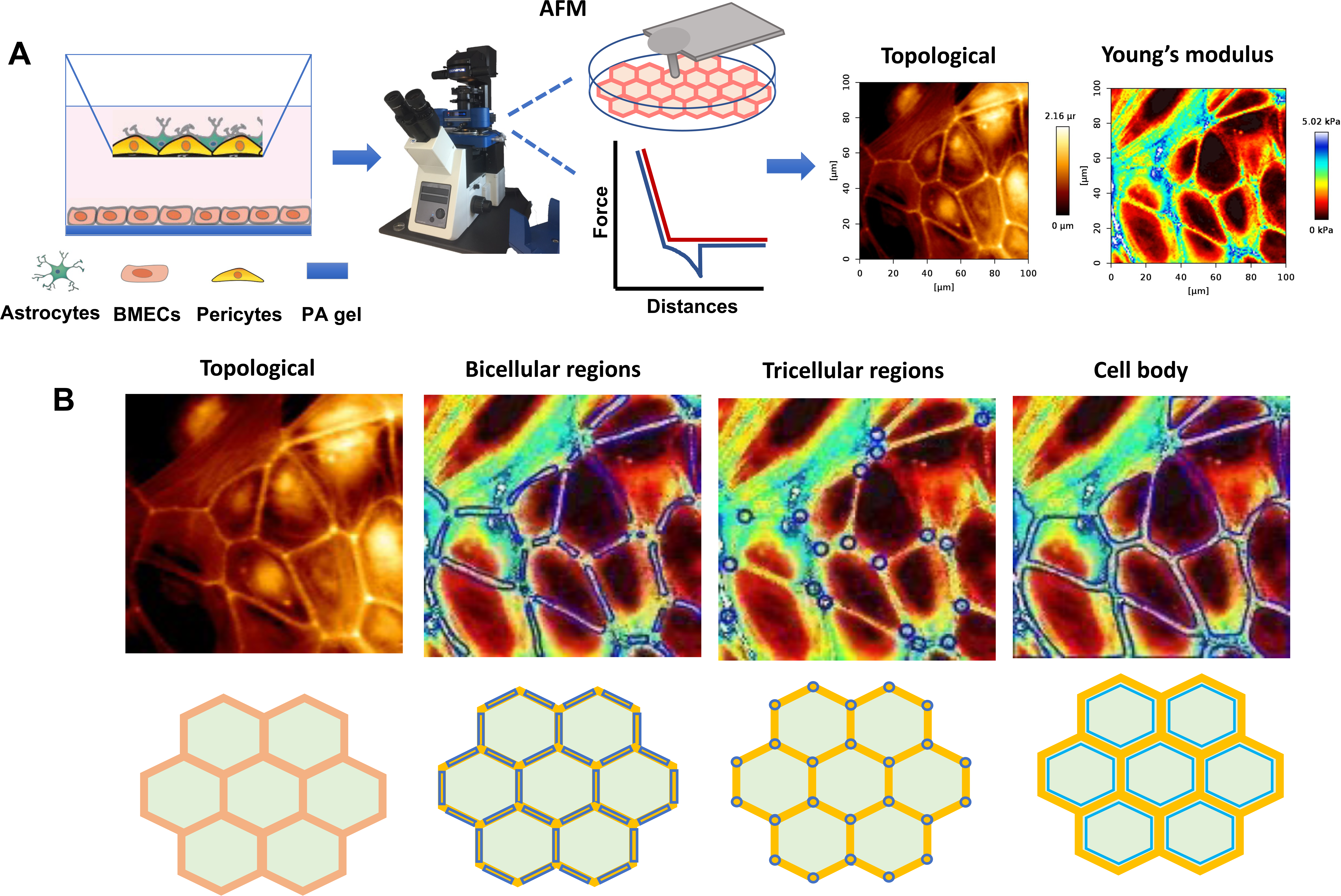
Schematic of BBB co-culture system and methodology for regional AFM-based mechanical analysis of iBMECs. (A) Experimental setup showing iBMECs cultured on PA hydrogels of tunable stiffness and co-cultured with astrocytes and pericytes. (B) Immunofluorescence confirms cell identity and phenotype: pericytes express PDGFRβ (green), astrocytes express GFAP (green), and iBMECs express the tight junction marker ZO-1 (red). Nuclei are stained with DAPI (blue). (C) AFM setup and representative maps showing topography and Young’s modulus of iBMECs. (D) Regional segmentation strategy used for analysis. MATLAB-based image processing defines bicellular junctions (between two cells), tricellular junctions (intersection of three cells), and cell body regions. All scale bars represent 50 μm.

To examine how matrix stiffness and BBB cell (astrocytes and pericytes) co-culture influence iBMEC mechanical behavior, we performed AFM on iBMECs on PA gels spanning a range of physiologic and pathologic stiffnesses (1, 2.5, 15, and 194 kPa), both alone and in co-culture with astrocytes and pericytes (iBAP). In iBMECs alone, increasing matrix stiffness led to pronounced changes in monolayer morphology and mechanical response. AFM modulus maps showed elevated stiffness, particularly along cell–cell junctions at stiffness 2.5 kPa and 15 kPa. However, on excessively stiff substrates (194 kPa), iBMECs exhibited a reduction in Young’s modulus (Figure 2A). In contrast, iBMECs in iBAP groups displayed consistently lower stiffness across all conditions. Notably, iBAP cultures also exhibited disrupted morphology and reduced stiffness on the 194 kPa substrate, similar to iBMECs alone (Figure 2B). These findings indicate that perivascular co-culture modulates iBMEC mechanical response to substrate stiffness.

**Figure 2.**
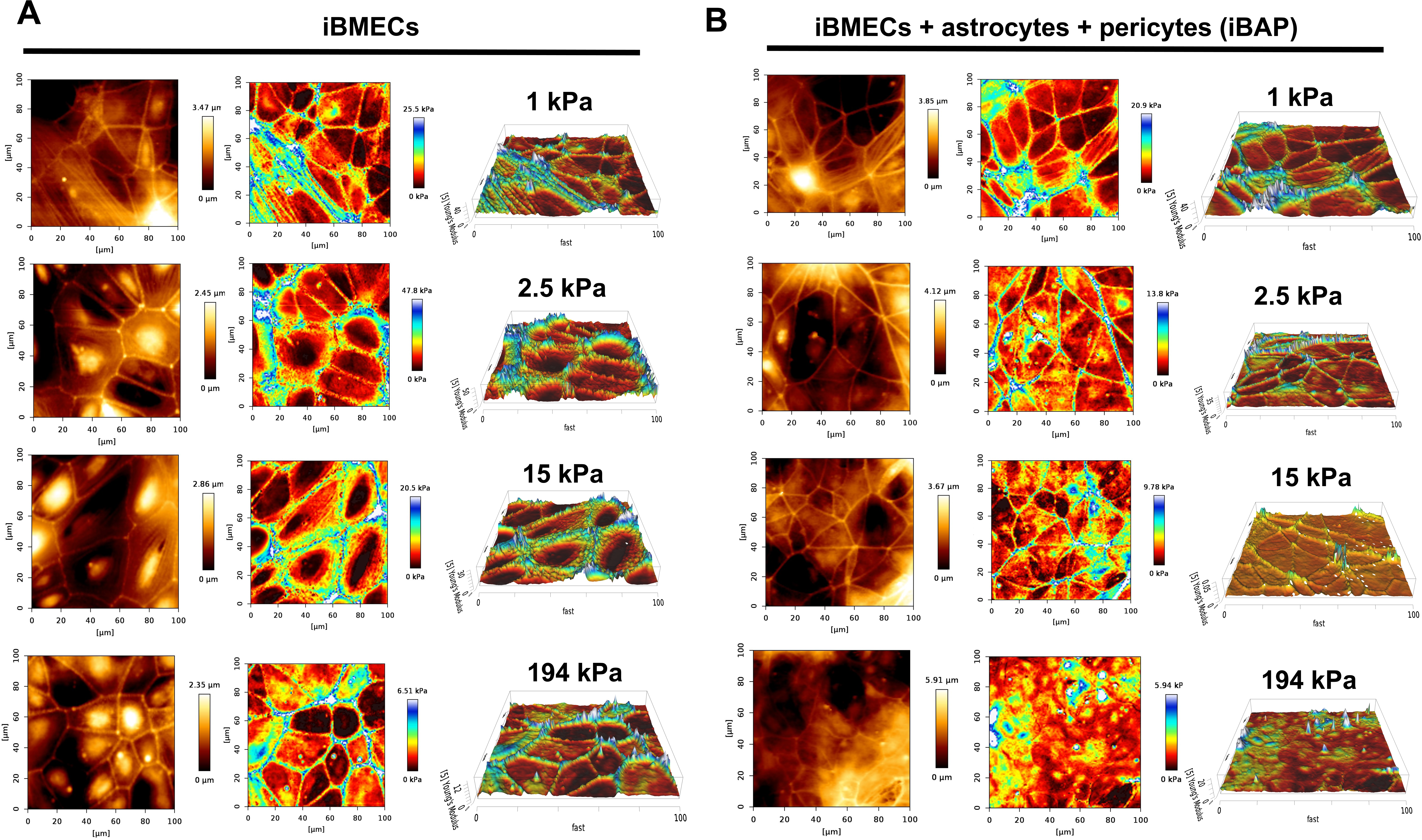
AFM characterization on monoculture and co-culture on PA hydrogels of increasing stiffness. (A) Representative AFM maps of IMR90-1 iBMEC monolayers cultured on 1, 2.5, 15, and 194 kPa substrates. From left to right: topography, Young’s modulus maps, and 3D surface reconstructions showing stiffness distribution. (B) Corresponding AFM data from IMR90-1 iBMECs co-cultured with astrocytes and pericytes (iBAP). Color bars represent height (µm) and Young’s modulus (kPa).

### Co-culture with astrocytes and pericytes reduces regional iBMEC stiffness and attenuates mechanical polarization across substrate stiffnesses

To evaluate subcellular mechanical differences in iBMECs and how they are influenced by substrate stiffness and BBB cell support, we measured the Young’s modulus of three distinct regions in iBMECs: tricellular junctions, bicellular junctions, and cell bodies. In Ibmec monocultures, cells cultured on physiologically relevant 2.5 kPa substrates exhibited significantly higher stiffness across all regions compared to those on excessively stiff 194 kPa substrates (Figure 3A). In addition, iBMECs on 2.5 and 15 kPa substrates showed a relatively broad distribution of stiffness values, with preserved differences between junctional and non-junctional regions (Figure S1). When iBMECs were co-cultured with astrocytes and pericytes (iBAP), overall stiffness decreased across all regions (Figure 3B). The narrowed and left-shifted distribution of relative frequency are further evidence of this effect (Figure S2). These results suggest reduced cytoskeletal tension and junctional stress in the presence of supporting BBB cell types. Interestingly, this decrease in stiffness was more pronounced on 2.5 kPa and 15 kPa substrates, particularly at tricellular and bicellular junctions, when compared to iBMECs cultured alone (Figure 3B). Notably, in contrast to iBMECs alone, iBAP cultures displayed significantly higher stiffness on the softest 1 kPa substrate relative to 194 kPa (Figure 3A). These results suggest a stiffness-dependent modulation of the mechanical phenotype by BBB cells.

**Figure 3.**
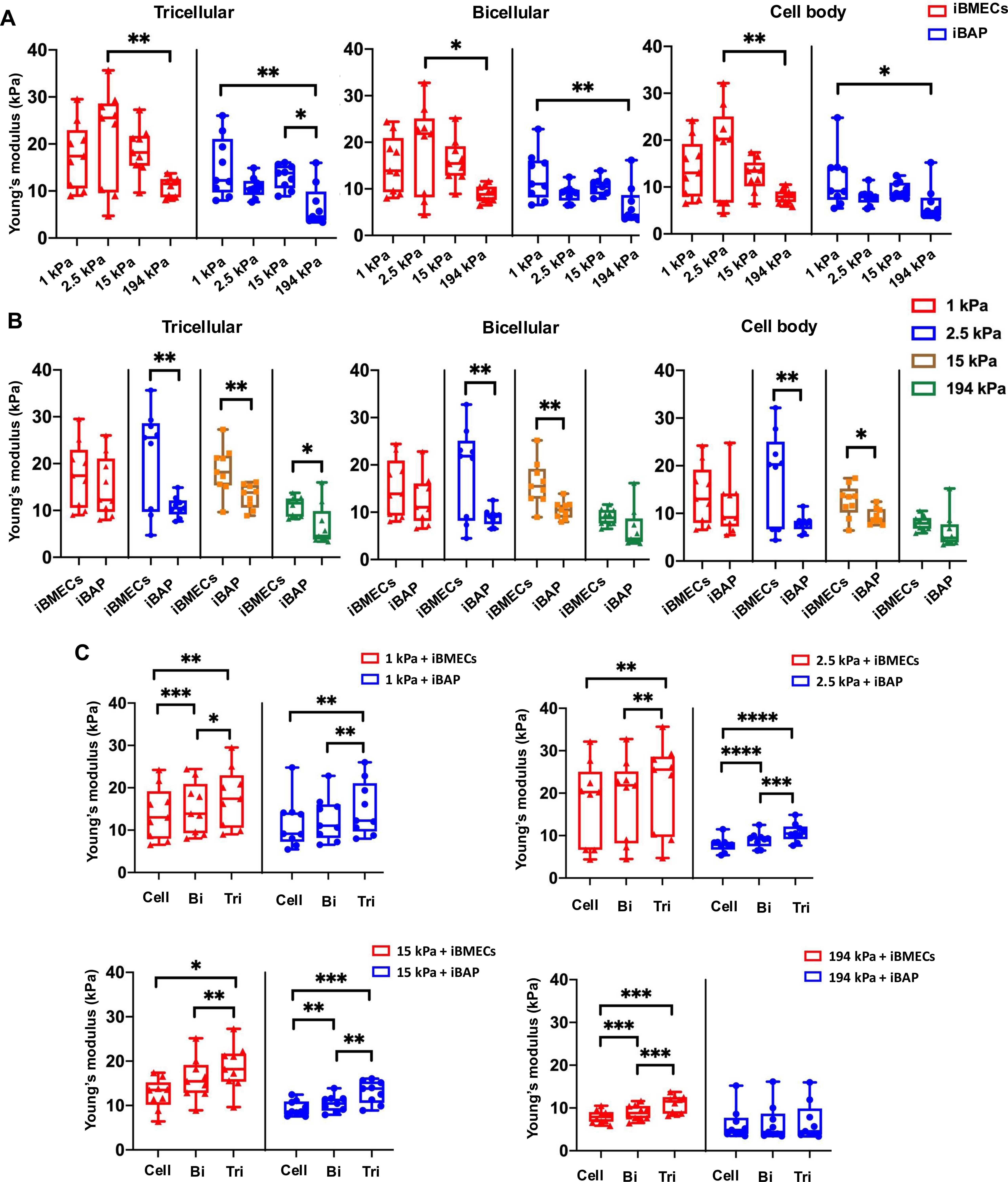
Quantification of regional stiffness in iBMEC monolayers across substrate stiffness and co-culture conditions. (A) Young’s modulus measurements at tricellular junctions, bicellular junctions, and the cell body in IMR90-1 iBMECs (red) and iBAP (blue) across four substrate stiffnesses (1, 2.5, 15, and 194 kPa). Statistical analysis was performed using one-way ANOVA with Tukey’s multiple comparisons post hoc test. Error bars represent standard deviation. *p<0.05, **p<0.01. (B) Direct comparison of iBMEC and iBAP stiffness within each substrate condition. Unpaired t-tests were used for statistical analysis. *p<0.05, **p<0.01. (C) Intracell comparisons of stiffness among tricellular, bicellular, and cell body regions within each group and matrix condition. Paired t-tests were used. *p<0.05, **p<0.01, ***p<0.001, ****p<0.0001.

A consistent pattern of regional mechanical polarization was observed in both monoculture and co-culture conditions: tricellular junctions are stiffer than bicellular junctions and cell bodies across most conditions (1, 2.5, and 15 kPa) (Figure 3C). However, this polarization was lost in iBAP cultures on 194 kPa substrates, where tricellular stiffness dropped sharply and regional differences became less distinct (Figure 3C). These findings indicate that soft (1 kPa and 2.5 kPa) and moderately stiff (15 kPa) substrates support the maintenance of junctional mechanical polarization, particularly at tricellular junctions, while supraphysiological stiffness (194 kPa) disrupts this organization, especially in the presence of BBB cells.

### MDA-231-BR cells alter mechanical architecture of iBMECs in iBAP co-culture

To evaluate how the regional mechanical modulation changes under simulated disease condition, we added MDA-MB-231-BR cells to iBAP culture and performed AFM to assess topography and regional stiffness of iBMECs (Figure 4A). In untreated iBAP cultures (independent experiments from Figure 2), topographical maps showed well-organized monolayers with robust junctional ridges, and stiffness maps revealed elevated Young’s modulus at intercellular junctions, particularly at tricellular points on soft gels (Figure 4B). Following MDA-MB-231-BR cell exposure, iBMECs displayed pronounced structural and mechanical disruption (Figure 4C). Topography maps revealed flatter, less organized surfaces, and elasticity maps showed marked reductions in stiffness across all regions. In several fields, junctional features were fragmented or absent, and Young’s modulus fell below 1 kPa. This widespread loss of mechanical integrity suggests that MDA-MB-231-BR cells compromise endothelial monolayers by disrupting junctional tension and weakening cytoskeletal reinforcement.

**Figure 4.**
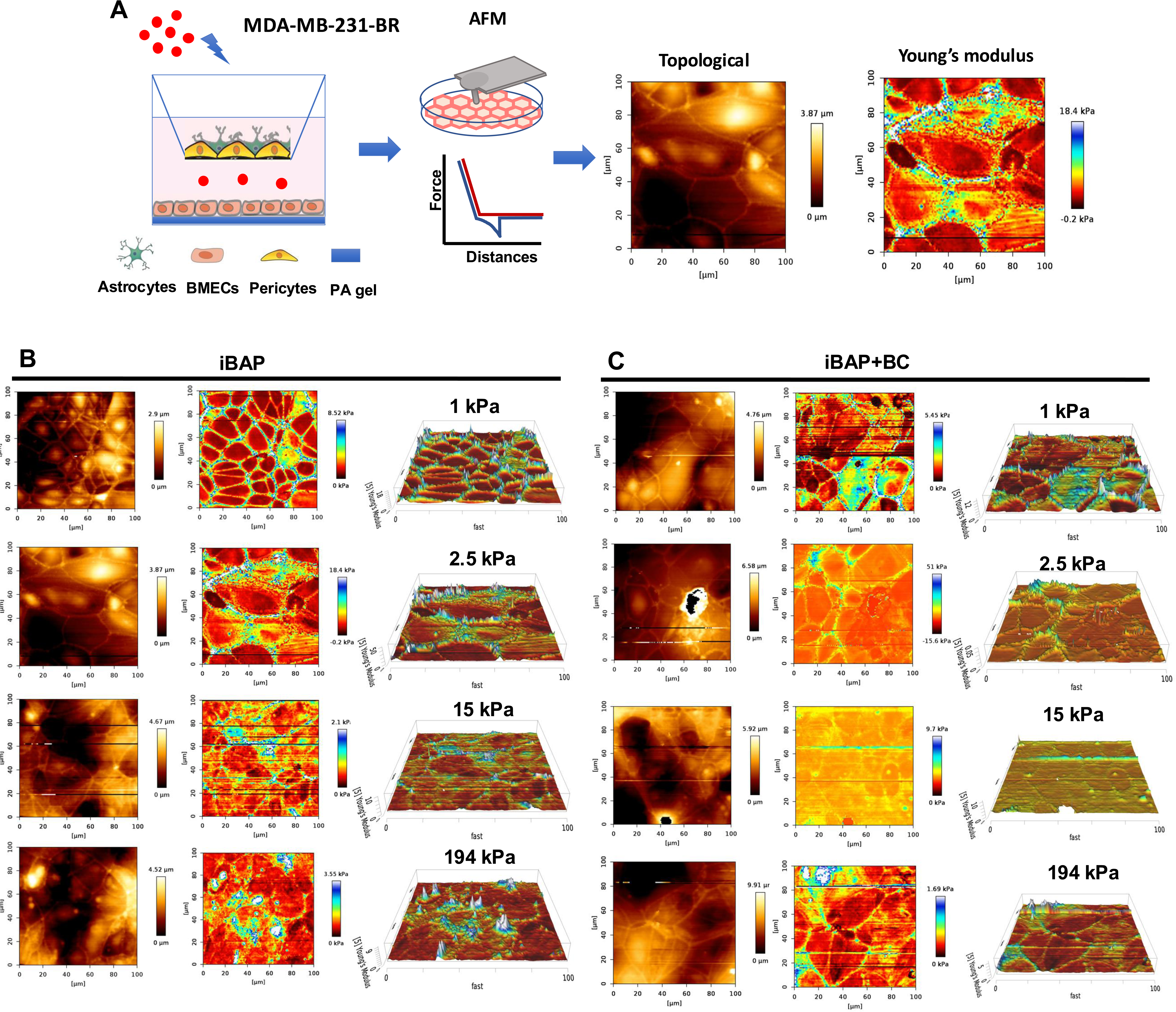
AFM characterization of iBMECs in BC co-culture model. (A) Schematic of experimental setup: MDA-231-BR cells were introduced to IMR90-1 iBAP monolayers cultured on polyacrylamide gels. AFM was performed to assess topography and Young’s modulus of the endothelial surface after BC exposure. (B) Representative AFM maps of iBAP cultures without cancer cells. Topography, Young’s modulus, and 3D stiffness reconstructions show intact monolayers with high stiffness at junctional regions, indicating mechanical polarization and barrier integrity. (C) Representative AFM maps of iBAP cultures after exposure to breast cancer cells (iBAP+BC). Color bars indicate height (µm) and Young’s modulus (kPa). Scale bars: 20 µm.

### MDA-231-BR cells disrupt junctional stiffness and preferentially transmigrate across iBMECs on softer hydrogels

To investigate how MDA-MB-231-BR cells influence BBB mechanics, we measured the regional Young’s modulus of iBAP monolayers before and after cancer cell exposure across substrates of varying stiffness. In co-culture conditions, overall stiffness across all regions did not significantly differ from iBAP controls (Figure 5A). However, in untreated iBAP cultures, junctional regions—particularly tricellular regions—were significantly stiffer than the cell bodies, especially on soft substrates (1 and 2.5 kPa), indicating preserved mechanical polarization (Figure 5B and Figure S3), consistent with our observations in Figure 3C. Exposure to MDA-MB-231-BR cells disrupted this mechanical organization, reducing stiffness of junctions at tricellular and bicellular regions and eliminating the distinct regional differences. The narrower, more left-skewed distributions of the relative frequency suggest that MDA-MB-231-BR cells reduces overall stiffness of iBMECs (Figure S4). These findings suggest that metastatic cancer cells actively remodel the endothelial monolayer, compromising its structural barrier integrity.

**Figure 5.**
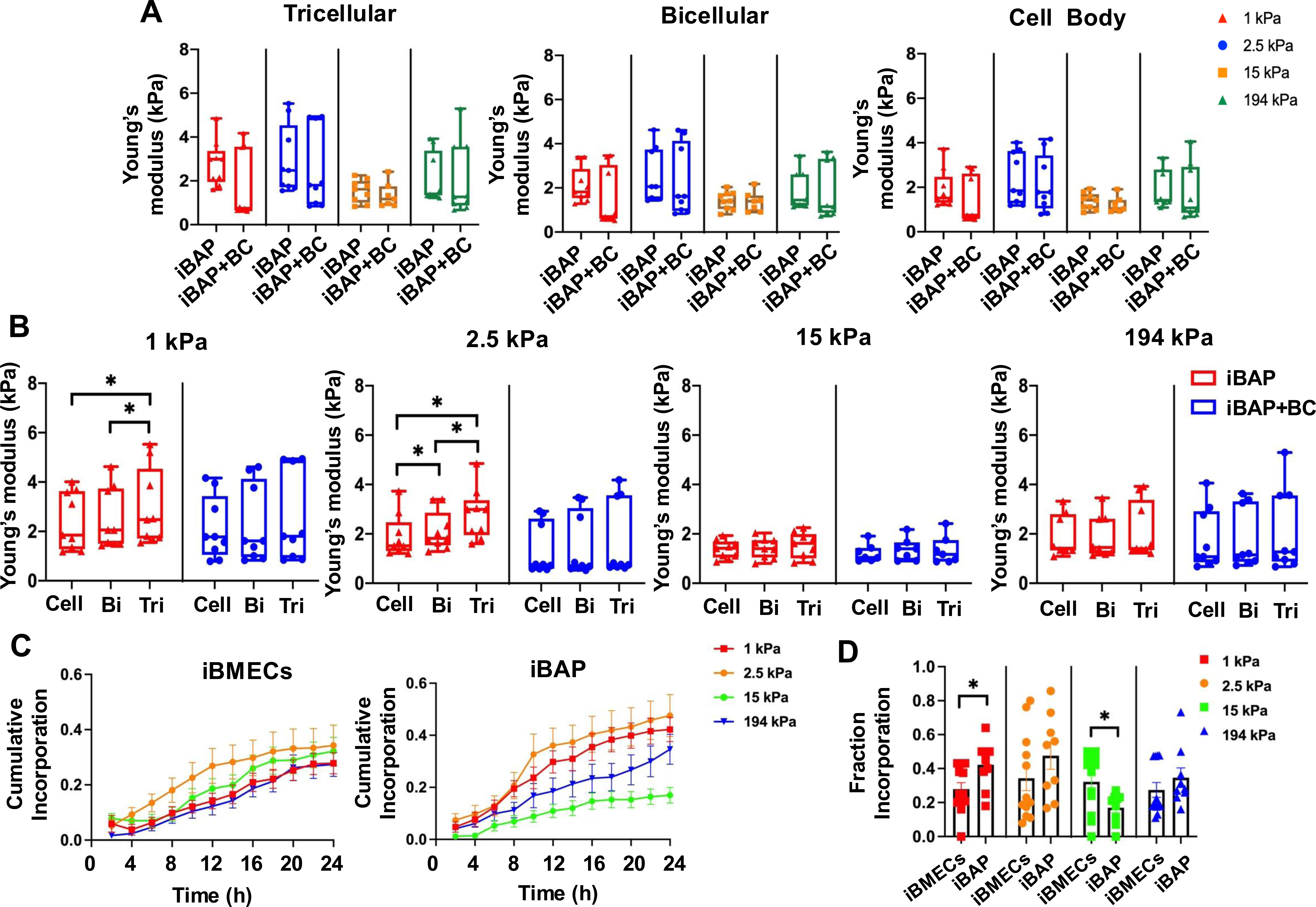
Comparison of regional stiffness in BC co-culture with BBB and BC incorporation. (A) Comparison of Young’s modulus across tricellular junctions, bicellular junctions, and cell body regions in IMR90-1 iBAP monolayers before and after BC cell exposure (iBAP+BC) at four substrate stiffnesses (1, 2.5, 15, and 194 kPa). (B) Within-condition analysis of regional stiffness in iBAP and iBAP+BC cultures. (C) Cumulative incorporation of BC cells over 24 hours in iBMEC and iBAP monolayers cultured on 1, 2.5, 15, or 194 kPa substrates. (D) Quantification of total BC cell incorporation fraction across conditions. Comparisons between iBMEC and iBAP were analyzed using unpaired t-tests. Error bars represent standard deviation. *p<0.05, **p<0.01.

To determine whether these mechanical changes facilitate transendothelial migration, we tracked cancer cell incorporation over 24 hours following co-culture with iBMECs or iBAP (Video S1). In iBMEC monocultures, MDA-MB-231-BR cells exhibited similar incorporation rates across all substrate stiffnesses. In contrast, iBAP co-cultures showed greater incorporation on softer matrices (1 and 2.5 kPa), with reduced entry on stiffer substrates (15 and 194 kPa) (Figure 5C). Statistically significant differences were observed in the 1 kPa and 15 kPa groups. These results indicate that while substrate stiffness alone has a minimal effect on cancer cell entry in monocultures, astrocytes and pericytes alter the mechanical barrier properties of the BBB. This renders the softer matrices more susceptible to invasion, whereas intermediate stiffness may promote preferential breaching of mechanically vulnerable sites.

### iBAP co-culture softens iBMECs through activation of mature and primed BBB transcriptomic signatures

AFM mapping results revealed that iBMECs co-cultured with astrocytes and pericytes (iBAP) condition exhibited reduced general cellular stiffness compared to monoculture across all ECM stiffness levels (1.5, 2.5, 15, and 194 kPa). This softening effect was most evident at cell junctions and borders, suggesting modulation of cortical tension and junctional mechanics. To investigate the molecular mechanisms underlying this biomechanical alteration, we performed bulk RNA sequencing on iBMECs in monoculture and iBAP co-culture. Principal component analysis (PCA) revealed clear segregation between the two conditions (Figure 6A), indicating that iBAP co-culture induces broad transcriptomic changes. Volcano plot analysis identified numerous differentially expressed genes (DEGs) between iBMEC and iBAP samples (Figure 6B), and heatmap clustering of the top 100 variable genes (Figure 6C) showed distinct grouping by co-culture status, independent of ECM stiffness, highlighting the dominant effect of cellular interactions over matrix mechanics. Gene set enrichment analysis (GSEA) revealed that iBAP co-culture significantly upregulated pathways involved in tight junction assembly, actin cytoskeleton remodeling, and cell adhesion signaling (Figure 6D). Additionally, pathways related to immune signaling (e.g., lymphocyte-mediated immunity), angiogenesis, and vascular development were enriched in the iBAP group, along with increased cell–cell adhesion signaling (Figure 6D). At the gene level, tight junction components (e.g. CLDN3, CLDN19, TJP3), extracellular matrix remodeling factors (e.g. MMP28, COL6A3, VWA1), and inflammatory mediators (e.g. IL1A, CXCL14) were significantly upregulated in iBAP co-culture, suggesting enhanced junctional regulation and cytoskeletal plasticity (Figure 6B). In contrast, a subset of genes associated with endothelial immaturity or angiogenic signaling, including EPHA7, SHC4, ETV5, and DACH1, were downregulated (Figure 6B), indicative of a transition toward a more stable, mature barrier phenotype. These transcriptomic signatures demonstrate that iBAP co-culture promotes a more mature and functionally primed BBB phenotype which weaken junctional and cell tension and account for the reduced cell stiffness.

**Figure 6.**
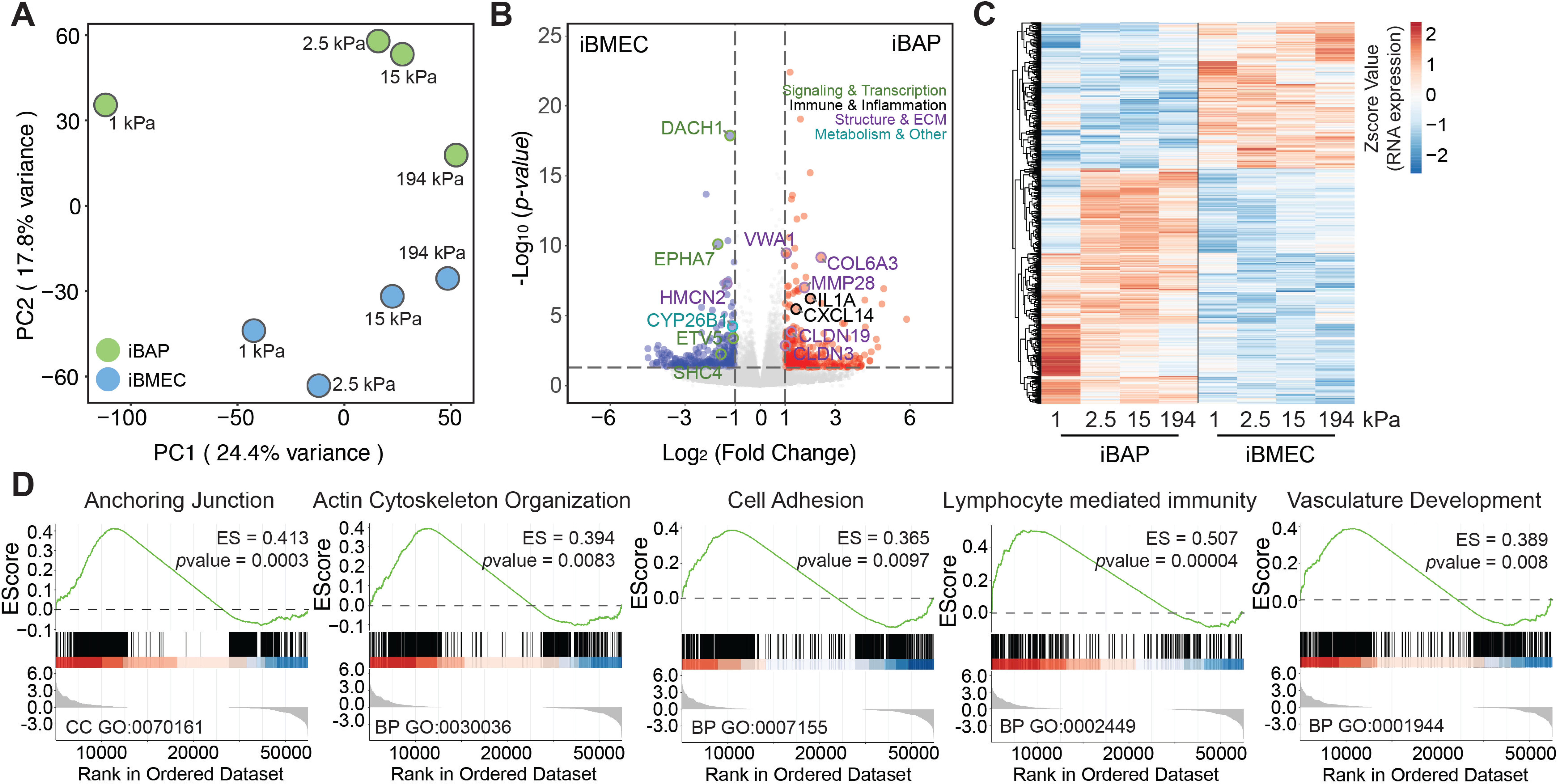
Transcriptomic analysis of stiffness-dependent regulation in iBMECs and iBAP co-cultures. (A) PCA showing distinct clustering of IMR90-1 iBMECs (blue) and iBMECs co-cultured with astrocytes and pericytes (iBAP, green) across different substrate stiffness (1, 2.5, 15, and 194 kPa. ^5^(B) Volcano plot comparing iBAP and iBMEC transcriptomes. (C) Heatmap of Z-score-normalized expression for stiffness-responsive genes. (D) GSEA identifying biological processes significantly enriched in iBAP relative to iBMECs, including anchoring junction organization, actin-cytoskeleton regulation, cell adhesion, lymphocyte-mediated immunity, and vasculature development.

### Transcriptomic profiling reveals stiffness-dependent transitions from junctional reinforcement to stress adaptation

Despite overall reductions in cell and matrix stiffness, stiffness mapping revealed that junctions at tricellular regions remained mechanically reinforced under soft ECM conditions (1 kPa and 2.5 kPa) in both iBMEC and iBAP cultures, maintaining higher stiffness than bicellular junctions or cell bodies. This pattern was absent on stiffer substrates (194 kPa). RNA-seq analysis provided molecular insight into these mechanical adaptations. Unsupervised clustering of differentially expressed genes across four stiffness conditions in both monoculture and co-culture models (Figure S5) identified 36 stiffness-associated gene expression patterns, with five major gene groups showing consistent trends across the stiffness gradient (Figure S6). Analysis of the gene core signature further revealed stiffness-dependent transcriptional profiles associated with distinct mechanical behaviors in iBMECs. On soft hydrogels, AFM confirmed that tricellular junctions exhibited higher local stiffness than neighboring bicellular regions, indicating active reinforcement at these vertices. Transcriptomic profiling supported this observation that cells under soft, brain-like stiffness, both groups exhibited enriched expression of genes associated mechanosensing and focal adhesion dynamics (e.g. PXN, TLN1, ACTN4, VCL, FLNA), mechanotransduction and angiogenic signaling (e.g. KDR, PIEZO1, TGFB2, VEGFA), and matrix remodeling (e.g. MMP9, TIMP3, SPARC, CTGF) were upregulated, together with adhesion and junctional components (e.g. JAM2, ESAM, ICAM1) (Figure S7). These gene profiles support an adaptive cytoskeletal state characterized by active force redistribution and dynamic junctional organization. In contrast, cells on stiff substrates (194 kPa) exhibited elevated expression of YAP1, TEAD1, and LATS1, indicating activation of canonical YAP/TAZ signaling under sustained tension, along with SRC, VCAM1, ANGPT, and COL6A3 (Figure S7), which are associated with ECM reinforcement and mechanical stress adaptation. Collectively, these results demonstrate a stiffness-dependent transcriptional transition from dynamic mechanoadaptation and junctional reinforcement on soft matrices to stress-driven, rigidity-sustaining programs on stiff environments.

## Discussion

BBB integrity arises from the coordinated interplay between ECM mechanics and neurovascular signaling, which together regulate cytoskeletal tension and force transmission within brain endothelial cells^20^. Yet, how these mechanical and cellular cues integrate to control local reinforcement remains poorly understood. Although vascular stiffening has been correlated with BBB dysfunction^20, 21, 22^, the mechanisms through which ECM stiffness and supporting cell signals modulate junctional mechanics in health and disease are still unclear. Here, we show that ECM stiffness, in concert with astrocytes and pericytes, governs the spatial distribution of mechanical forces across brain endothelial subcellular regions, with tricellular regions functioning as specialized vertices that serve as hubs for stress concentration and cytoskeletal anchoring.

AFM revealed that iBMECs display regional mechanical polarization, with tricellular junction regions exhibiting higher stiffness than bicellular junctions and cell bodies, particularly on compliant hydrogels (1–15 kPa) indicating active redistribution of mechanical reinforcement rather than uniform contractility. In contrast, this polarization was lost under supraphysiological stiffness (194 kPa), where overall cell stiffness decreased and regional distinctions were eliminated. Together with our previous findings that junctions at tricellular regions act as local regulators of permeability^6^, these results identify tricellular vertices as structural and functional mechanical valves where endothelial tension and junctional architecture converge to maintain barrier stability.

Astrocytes and pericytes stabilized endothelial junctions and supported barrier function through both biochemical and mechanical regulation^23, 24, 25, 26^. Co-culture with these supporting cells reduced global stiffness while preserving regional polarization, demonstrating that BBB-associated cells buffer cytoskeletal tension while maintaining subcellular organization. This mechanical heterogeneity suggests that soft ECM environments promote adaptive cytoskeletal remodeling and localized reinforcement at multicellular contacts. RNA-seq analysis further confirmed that co-cultured iBMECs exhibit a transcriptionally primed BBB state characterized by dynamic and flexible junctional organization under compliant condition, while cells on stiff substrates lead to strong contractile forces that resist mechanical forces and less flexibility ^27, 28,29^. Together, these findings identify homeostatic compliance as a spatially organized mechanical regime required for healthy BBB function, reflecting a transition from stress-fiber–mediated rigidity to localized, junction-centered force regulation.

The differential stiffness affects cancer cell brain invasions^30^. In our studies, exposure to metastatic breast cancer–derived factors profoundly disrupted BBB mechanics, abolishing tricellular polarization and suppressing stiffness across all regions. These findings indicate that metastatic breast cancer cells compromise BBB integrity by altering cytoskeletal organization and disrupting junctional mechanotransduction, potentially facilitating tumor cell extravasation and metastasis. Notably, breast cancer cells preferentially integrated into soft (1 kPa) hydrogels, where junctions were more flexible and responsive to external cues. These observations indicate that compliant junctions provide enhanced mechanical plasticity that facilitates invasion, in contrast to the rigid, stress-fiber–dominated resistance characteristic of stiffer substrates. Together, these observations highlight the critical role of spatially organized mechanics in BBB function, revealing how both physiological and pathological changes in the microenvironment reshape barrier properties.

Collectively, our findings establish a spatially resolved framework for BBB mechanobiology. Physiological stiffness and BBB cell interactions maintain regional mechanical heterogeneity and transcriptional balance. Tricellular junction regions emerge as key biomechanical and regulatory nodes where physical and molecular mechanisms intersect to govern barrier function.

Targeting this local mechanoadaptive machinery may offer new therapeutic opportunities to preserve BBB integrity or selectively modulate permeability in neurological disease and metastasis.

## Methods

### iPSC culture and BMECs differentiation

IMR90-1 and DF19-1-19T.H (DF19) human induced pluripotent stem cells (iPSCs) (WiCell) were maintained on Matrigel (Corning)-coated 6-well plates in mTeSR medium (STEMCELL Technologies), as previously described. For differentiation into iBMECs, iPSCs were dissociated using Accutase (Thermo Fisher Scientific) and seeded onto Matrigel-coated 6-well plates at a density of 1.0–1.25 × 10L cells/well in Essential 8 medium supplemented with 10 µM Y27632 (R&D Systems). After 24 hours, the medium was switched to Essential 6 medium (Thermo Fisher Scientific) to initiate differentiation (designated as day 0). On day 4, the medium was replaced with endothelial cell (EC) medium. EC medium is composed of human endothelial serum-free medium (Thermo Fisher Scientific) supplemented with 1% platelet-poor plasma-derived serum (Thermo Fisher Scientific), 20 ng/mL basic fibroblast growth factor (bFGF, PeproTech), and 10 mM ascorbic acid (Sigma-Aldrich). On day 6, cells were dissociated into single cells using Accutase and seeded onto polyacrylamide (PA) hydrogels coated with human placenta-derived type IV collagen (400 µg/mL, Sigma-Aldrich) and human plasma-derived fibronectin (100 µg/mL, Sigma-Aldrich). On day 7 (one day post-subculture), the culture medium was replaced with EC medium lacking RA and bFGF for maintenance.

### Pericyte differentiation

Pericytes was differentiated as previously described^31^. Briefly, iPSCs were dissociated with Accutase and seeded onto Matrigel-coated 6-well plates in mTeSR medium (STEMCELL) containing 10 μM Y-27632 (R&D System). After 24 h, medium was changed into pericyte differentiation medium (Day 0). Pericyte differentiation medium is composed of E6 medium (Thermo Fisher Scientific) supplemented with 1μM CHIR99021, 10 μM SB431542 (STEMCELL), 10 ng/ml bFGF (Peprotech), and 1μM dorsomorphin (Tocris). Cells were cultured in pericyte differentiation medium for 15 days. Cells were dissociated and replated in E6 medium supplemented with 10% fetal bovine serum (FBS) (Thermo Fisher Scientific) for 10 days. After that, cells were maintained in E6 medium containing 10% FBS until they were co-cultured.

### Astrocyte differentiation

Astrocytes were differentiated as previously described^23^. Briefly, iPSCs were dissociated with Accutase and seeded onto Matrigel-coated 6-well plates at 2×10^5^ cells/cm^2^ in mTeSR medium (STEMCELL) containing 10 μM Y-27632 (R&D System). 24 h after seeding, medium was changed to E6 medium (Thermo Fisher) supplemented with 10 μM SB431542 (Tocris) and 1 μM dorsomorphin dihydrochloride (Tocris) (Day 0). The medium was changed daily for 6 days. On day 6, cells were detached and were seeded to a Matrigel-coated plate with E6 medium supplemented with 10 ng/mL epidermal growth factor (EGF) (Peprotech), 10 ng/mL ciliary neurotrophic factor (CNTF) (Peprotech), and 10 μM Y-27632 (R&D System). After 24h, the medium was changed to E6 medium containing 10 ng/mL CNTF and 10 ng/mL EGF. The medium was changed every 3 days for 30 days. After that, cells were maintained in E6 medium containing 10 ng/mL CNTF and 10 ng/mL EGF until they were co-cultured.

### Breast cancer (BC) cell culture

The brain-seeking clones of human breast adenocarcinoma cells (MDA-MB-231-BR) were cultured in breast cancer medium, which includes Dulbecco’s modified eagle’s medium with high glucose and L-glutamine supplemented with 10% fetal bovine serum (FBS) and 1% penicillin/ streptomycin (Thermo Fisher Scientific).

### Polyacrylamide (PA) hydrogel preparation

PA hydrogels were polymerized on glass-bottom 35 mm cell culture dishes (WPI). To generate hydrogels with defined stiffness values, acrylamide and bis-acrylamide (Bio-Rad) were mixed at the following concentrations: 3% acrylamide + 0.2% bis (1 kPa), 7.5% acrylamide + 0.075% bis (2.5 kPa), 8% acrylamide + 0.2% bis (15 kPa), and 15% acrylamide + 1.2% bis (194 kPa). The Young’s modulus of each hydrogel formulation was previously validated using atomic force microscopy (AFM)^6, 13^, as described in our prior publication. Following polymerization, the hydrogel surfaces were functionalized with sulfo-SANPAH (Thermo Fisher Scientific) and activated under UV light. Gels were then washed three times with PBS and coated overnight at 4°C with 400 µg/mL collagen type IV and 100 µg/mL fibronectin. Prior to cell seeding, hydrogels were washed twice with PBS and pre-equilibrated in culture medium for 10 minutes. On differentiation day 6, BMECs were seeded onto the PA hydrogels at a density of 2 × 10L cells per dish for AFM measurements.

### Co-culture experiment

On differentiation day 6, 1.5 × 10L iBMECs were seeded onto PA hydrogels in 35 mm cell culture dishes (WPI). On the same day, human iPSC-derived astrocytes and pericytes were seeded onto 24 mm Transwell inserts at densities of 2 × 10L and 3 × 10L cells per insert, respectively, using a 1:1 mixture of astrocyte and pericyte culture media. After 24 hours (day 7), the Transwell inserts containing astrocytes and pericytes were transferred onto the iBMEC-seeded hydrogels to establish the co-culture system (designated as iBAP). The medium was then changed to endothelial cell EC medium without RA or bFGF for continued co-culture. On day 9 (48 hours after initiating co-culture), atomic force microscopy (AFM) measurements were performed on the iBMECs derived from both IMR90-1 and DF19. For cancer cell co-culture, 2 × 10L MDA-MB-231-BR cells were added directly to the bottom compartment of the iBAP system derived from IMR90 on day 9. After 24 hours of co-culture (day 10), iBMECs were subjected to AFM scanning.

### Cancer cell incorporation assay and time-lapse imaging

To assess cancer cell incorporation into the iBAP, the co-culture medium was first collected on day 9 of differentiation. iBMECs were gently washed twice with DPBS to remove unattached cells or debris. The collected medium was then filtered through a 0.22 μm sterile filter and returned to the same well to maintain consistent culture conditions. Next, 2 × 10L MDA-MB-231-BR cells were added to the bottom compartment of the iBAP system. Immediately following cancer cell seeding, the cultures were transferred to a live-cell imaging chamber maintained at 37°C, 5% CO₂, and controlled humidity. Time-lapse imaging was performed using an automated inverted fluorescence microscope (IX83, Olympus) equipped with a motorized stage and environmental control. Images were acquired every 10 minutes for a total duration of 24 hours. Three representative fields of view were selected per well for continuous imaging. After image acquisition, time-lapse sequences were analyzed using ImageJ (NIH) to quantify the cumulative incorporation of cancer cells over time. Incorporation was defined based on cancer cell spreading into the iBMEC monolayer^32^. Quantification was performed across three independent biological replicates.

### Atomic force microscopy

All AFM experiments were conducted using a NanoWizard 4a BioScience AFM system (JPK Instruments AG) equipped with SAA-SPH-1UM cantilevers (Bruker) with nominal spring constants ranging from 0.15 to 0.32 N/m. The actual spring constant of each cantilever was calibrated prior to each experiment using the thermal tune method and was consistent with the manufacturer’s specifications. Mechanical measurements were performed in Quantitative Imaging (QI) mode. For each experimental condition, three distinct 100 × 100 μm regions were imaged across three separate cell areas. Each region was captured at a resolution of 128 × 128 pixels. Force–distance curves were acquired using the JPK AFM software and subsequently processed and analyzed with JPK image processing software (JPK Instruments AG). For each substrate stiffness condition, three independent regions were imaged per sample. All experiments were conducted in three biological replicates.

### AFM image analysis

To quantitatively analyze the stiffness of different regions within the iBMECs, a MATLAB-based program was developed to analyze the AFM images. Using the MATLAB tool, Young’s Modulus data obtained from AFM was associated on a per-pixel basis with the AFM images and distinct regions of the cells were waypointed and analyzed. The program first corrects the Young’s Modulus data obtained from AFM and assigns each data point to the corresponding pixel. Once the Young’s Modulus data has been assigned to each pixel, the user is prompted to select the AFM image for analysis. First, the topological AFM image of the cells is chosen for waypointing. After selecting the first region of interest (ROI) on the topological image using MATLAB’s freehand object, the Young’s Modulus image map is selected, and the MATLAB program overlays the ROI selection and retrieves the data from the pixels selected within the ROI. Three different regions of the cells were considered for analysis from AFM. The cell body constitutes a major portion of the images. The bicellular junction covers the junctions neighboring two adjacent cells. For the analysis of the bicellular junctions, the ROI selection was at most 6 pixels in width. Any junction adjacent to three or more cells constituted the tricellular junctions. Circles with 6-pixel diameters were used to select for tricellular junctions. The topological image was used to identify these distinct regions.

### Immunochemistry

Immunocytochemistry was conducted on iBMECs, pericytes and astrocytes. Cells were fixed in 4% paraformaldehyde (Thermo Fisher) for 10 min. Cells were then treated with 0.25% Triton-X100 (Sigma) for 10 min. Cells were washed twice with PBS and blocked in 3% goat serum (Abcam) for 1h at room temperature. Primary antibodies were diluted in 3% goat serum, and cells were incubated in the primary antibodies at 4°C overnight. The next day, cells were washed three times with PBS. After that, cells were incubated in secondary antibody for 1h at room temperature. Cells were then washed with PBS three times, followed by treatment with Hoechst (Thermo Fisher Scientific) for 5 min. Cells were rinsed three times with PBS and then mounted. Cells were visualized and images were taken on the FV3000 Laser Scanning Confocal Microscope (Olympus). Antibody sources and dilutions are provided in Table S1.

### RNA isolation and sequencing

IMR90-1 derived iBMECs were harvested from both monolayer cultures and iBAP groups. Total RNA was extracted using the RNeasy Mini Kit (Qiagen) according to the manufacturer’s protocol. RNA quantity and purity were assessed with a NanoDrop spectrophotometer. RNA sequencing was performed by Novogene. Briefly, Messenger RNA was enriched from total RNA using poly-T oligo-attached magnetic beads. The mRNA was then fragmented and reverse transcribed into first-strand cDNA using random hexamer primers. Second-strand cDNA synthesis was performed using either dUTP (for directional libraries) or dTTP (for non-directional libraries). For both library types, subsequent steps included end repair, A-tailing, adapter ligation, size selection, PCR amplification, and purification. Directional libraries underwent an additional USER enzyme digestion step prior to amplification. Library quality and concentration were assessed using Qubit fluorometry, real-time PCR, and bioanalyzer for size distribution. Final libraries were pooled based on concentration and sequencing depth requirements. Cluster generation and paired-end sequencing were performed on an Illumina platform following the manufacturer’s protocols.

### Bulk RNA-seq Data Processing and Differential Expression

Raw FASTQ files were quality-filtered and adapter-trimmed, then aligned to the GRCh38 human genome reference. Gene-level counts were obtained and imported into R (v4.0). Low-abundance genes were removed, and data were normalized using DESeq2 with variance-stabilizing transformation (VST) for downstream visualization. Sample-level quality control included library size inspection, count distribution assessment, and outlier detection by PCA and sample distance heatmaps. Differential expression analysis was performed with a design matrix accounting for co-culture status and substrate stiffness. Wald statistics were used to estimate fold-changes, and Benjamini–Hochberg correction was applied to control false-discovery rate. Differentially expressed genes (DEGs) were defined as pvalue < 0.05 with directionality determined by log₂ fold-change. PCA plots, volcano plots, and hierarchical clustering heatmaps were generated in R to visualize global transcriptomic separation between groups.

### Gene Set Enrichment and Pathway Analysis

Rank-ordered log₂ fold-change values were used for pre-ranked GSEA using clusterProfiler with the fgsea algorithm. Enrichment was performed against Gene Ontology (BP, MF, CC) and Reactome pathway databases, with minimum 10 and maximum 2,000 genes per gene set. Enrichment significance was defined at pvalue < 0.05, and enrichment scores (ES) were used for ranking biological pathways. Representative GSEA running-score plots were generated for key pathways. Significantly enriched GO terms from each stiffness condition were aggregated, and a unified −log10(p value) enrichment matrix was constructed. Top pathways per group were identified based on nominal p-value rank. Pathway-by-condition matrices were clustered and visualized using pheatmap, with separate heatmaps for BP/MF/CC to distinguish biological, molecular, and cellular signatures. Processed enrichment tables and pathway matrices were exported to Excel for reproducibility.

### Variability-Driven Gene Clustering and Trajectory Patterns

Protein-coding genes (GENCODE v19 annotation) were ranked by median absolute deviation (MAD) across stiffness conditions. Highly variable genes were z-score–normalized and clustered using Gaussian mixture modeling (mclust). The optimal cluster number was selected via Bayesian Information Criterion (BIC), and the EVI covariance model was used, yielding 36 gene clusters. Cluster-wise expression trajectories across stiffness levels were min-max scaled, plotted with mean ± s.d., and visualized with ggplot2. Cluster membership tables were annotated and exported; identical processing was applied to the monoculture dataset to enable direct comparison. Analyses were performed in R (v4.0) using DESeq2, tidyverse, mclust, clusterProfiler, fgsea, pheatmap, ggplot2, and standard annotation packages.

### Statistical analysis

GraphPad Prism 10 was used for all statistical analysis and graph generation. For statistical analysis, a D’Agostino-Pearson normality test was performed to identify the normality of the data. If the data was normal, a one-way ANOVA with Tukey’s multiple comparison post hoc testing was used for analysis. The non-parametric Kruskal–Wallis ANOVA test with Dunn’s multiple comparison post hoc testing was used for the data sets that were not normally distributed. Comparisons between two groups were performed using two-tailed Student’s t-test. The specific statistical test used for each experiment is indicated in the corresponding figure legends. Statistical significance was indicated as *p < 0.05, **p < 0.01, ***p < 0.001, ****p < 0.0001. Errors bars represent the standard deviation of the mean as noted in the figure caption. All data represent are from three biological replicates. All iBMEC and iBAP experiments were performed using two independent human iPSC lines (IMR90 and DF19) to ensure reproducibility. Cancer cell incorporation assays and bulk RNA sequencing were conducted using IMR90-derived co-cultures due to their experimental consistency.

## Authors’ contributions

LY and KMS designed the research and wrote the manuscript. LY performed cell culture, immunostaining, AFM, and all other experiments. UG performed MATLAB analysis. HW performed RNA-seq analysis. All authors read and approved the final manuscript.

## Supporting information

Cancer cell incorporation

## Reference

1. Ferro MP, Heilshorn SC, Owens RM. Materials for blood brain barrier modeling in vitro. Mater Sci Eng R Rep 2020, 140.

2. Ryu Y, Iwashita M, Lee W, Uchimura K, Kosodo Y. A Shift in Tissue Stiffness During Hippocampal Maturation Correlates to the Pattern of Neurogenesis and Composition of the Extracellular Matrix. Front Aging Neurosci 2021, 13: 709620.

3. Streitberger KJ, Sack I, Krefting D, Pfuller C, Braun J, Paul F, et al. Brain viscoelasticity alteration in chronic-progressive multiple sclerosis. PLoS One 2012, 7(1): e29888.

4. Murphy MC, Huston J, 3rd, Jack CR, Jr., Glaser KJ, Manduca A, Felmlee JP, et al. Decreased brain stiffness in Alzheimer’s disease determined by magnetic resonance elastography. J Magn Reson Imaging 2011, 34(3): 494–498.

5. Chauvet D, Imbault M, Capelle L, Demene C, Mossad M, Karachi C, et al. In Vivo Measurement of Brain Tumor Elasticity Using Intraoperative Shear Wave Elastography. Ultraschall Med 2016, 37(6): 584–590.

6. Yan L, Dwiggins CW, Moriarty RA, Jung JW, Gupta U, Brandon KD, et al. Matrix stiffness regulates the tight junction phenotypes and local barrier properties in tricellular regions in an iPSC-derived BBB model. Acta Biomater 2023.

7. Katt ME, Linville RM, Mayo LN, Xu ZS, Searson PC. Functional brain-specific microvessels from iPSC-derived human brain microvascular endothelial cells: the role of matrix composition on monolayer formation. Fluids Barriers CNS 2018, 15(1): 7.

8. Sharif Y, Jumah F, Coplan L, Krosser A, Sharif K, Tubbs RS. Blood brain barrier: A review of its anatomy and physiology in health and disease. Clin Anat 2018, 31(6): 812–823.

9. Villabona-Rueda A, Erice C, Pardo CA, Stins MF. The Evolving Concept of the Blood Brain Barrier (BBB): From a Single Static Barrier to a Heterogeneous and Dynamic Relay Center. Front Cell Neurosci 2019, 13: 405.

10. Yan L, Moriarty RA, Stroka KM. Recent progress and new challenges in modeling of human pluripotent stem cell-derived blood-brain barrier. Theranostics 2021, 11(20): 10148–10170.

11. Bosworth AM, Kim H, O’Grady KP, Richter I, Lee L, O’Grady BJ, et al. Influence of Substrate Stiffness on Barrier Function in an iPSC-Derived In Vitro Blood-Brain Barrier Model. Cell Mol Bioeng 2022, 15(1): 31–42.

12. Haas AJ, Zihni C, Ruppel A, Hartmann C, Ebnet K, Tada M, et al. Interplay between Extracellular Matrix Stiffness and JAM-A Regulates Mechanical Load on ZO-1 and Tight Junction Assembly. Cell Rep 2020, 32(3): 107924.

13. Gray KM, Katz DB, Brown EG, Stroka KM. Quantitative Phenotyping of Cell-Cell Junctions to Evaluate ZO-1 Presentation in Brain Endothelial Cells. Ann Biomed Eng 2019, 47(7): 1675–1687.

14. Stroka KM, Aranda-Espinoza H. Endothelial cell substrate stiffness influences neutrophil transmigration via myosin light chain kinase-dependent cell contraction. Blood 2011, 118(6): 1632–1640.

15. Stroka KM, Levitan I, Aranda-Espinoza H. OxLDL and substrate stiffness promote neutrophil transmigration by enhanced endothelial cell contractility and ICAM-1. J Biomech 2012, 45(10): 1828–1834.

16. Gray KM, Stroka KM. Vascular endothelial cell mechanosensing: New insights gained from biomimetic microfluidic models. Semin Cell Dev Biol 2017, 71: 106–117.

17. Stroka KM, Aranda-Espinoza H. A biophysical view of the interplay between mechanical forces and signaling pathways during transendothelial cell migration. FEBS J 2010, 277(5): 1145–1158.

18. Rheinlaender J, Dimitracopoulos A, Wallmeyer B, Kronenberg NM, Chalut KJ, Gather MC, et al. Cortical cell stiffness is independent of substrate mechanics. Nat Mater 2020, 19(9): 1019–1025.

19. Bastounis EE, Yeh YT, Theriot JA. Subendothelial stiffness alters endothelial cell traction force generation while exerting a minimal effect on the transcriptome. Sci Rep 2019, 9(1): 18209.

20. Konig S, Jayarajan V, Wray S, Kamm R, Moeendarbary E. Mechanobiology of the blood-brain barrier during development, disease and ageing. Nat Commun 2025, 16(1): 7233.

21. Ling YH, Pan LH, Lin CH, Wang YF, Wu CH, Lirng JF, et al. Association of Increased Central Arterial Stiffness With BBB Disruption in Patients With Reversible Cerebral Vasoconstriction Syndrome. Neurology 2025, 105(10): e214318.

22. Reeve EH, Barnes JN, Moir ME, Walker AE. Impact of arterial stiffness on cerebrovascular function: a review of evidence from humans and preclincal models. Am J Physiol Heart Circ Physiol 2024, 326(3): H689–H704.

23. Neal EH, Marinelli NA, Shi Y, McClatchey PM, Balotin KM, Gullett DR, et al. A Simplified, Fully Defined Differentiation Scheme for Producing Blood-Brain Barrier Endothelial Cells from Human iPSCs. Stem Cell Reports 2019, 12(6): 1380–1388.

24. Canfield SG, Stebbins MJ, Morales BS, Asai SW, Vatine GD, Svendsen CN, et al. An isogenic blood-brain barrier model comprising brain endothelial cells, astrocytes, and neurons derived from human induced pluripotent stem cells. J Neurochem 2017, 140(6): 874–888.

25. Canfield SG, Stebbins MJ, Faubion MG, Gastfriend BD, Palecek SP, Shusta EV. An isogenic neurovascular unit model comprised of human induced pluripotent stem cell-derived brain microvascular endothelial cells, pericytes, astrocytes, and neurons. Fluids Barriers CNS 2019, 16(1): 25.

26. Jamieson JJ, Linville RM, Ding YY, Gerecht S, Searson PC. Role of iPSC-derived pericytes on barrier function of iPSC-derived brain microvascular endothelial cells in 2D and 3D. Fluids Barriers CNS 2019, 16(1): 15.

27. Sri-Ranjan K, Sanchez-Alonso JL, Swiatlowska P, Rothery S, Novak P, Gerlach S, et al. Intrinsic cell rheology drives junction maturation. Nat Commun 2022, 13(1): 4832.

28. Ahmed DW, Tan ML, Liu Y, Gabbard J, Gao E, Roy A, et al. Local photocrosslinking of native tissue matrix regulates lung epithelial cell mechanosensing and function. Nat Mater 2025, 24(11): 1812–1825.

29. Yates AK, Murray H, Kjar A, Chavarria D, Masters H, Kim H, et al. Substrate stiffness and shear stress collectively regulate the inflammatory phenotype in cultured human brain microvascular endothelial cells. Fluids Barriers CNS 2025, 22(1): 73.

30. Uroz M, Stoddard AE, Sutherland BP, Courbot O, Oria R, Li L, et al. Differential stiffness between brain vasculature and parenchyma promotes metastatic infiltration through vessel co-option. Nat Cell Biol 2024, 26(12): 2144–2153.

31. Stebbins MJ, Gastfriend BD, Canfield SG, Lee MS, Richards D, Faubion MG, et al. Human pluripotent stem cell-derived brain pericyte-like cells induce blood-brain barrier properties. Sci Adv 2019, 5(3): eaau7375.

32. Hamilla SM, Stroka KM, Aranda-Espinoza H. VE-cadherin-independent cancer cell incorporation into the vascular endothelium precedes transmigration. PLoS One 2014, 9(10): e109748.

