## Supplemental figure legends and table for "Substrate stiffness and cellular microenvironment regulate cell and junction mechanics in iPSC-derived brain microvascular endothelial cells"

**Figure S1. Substrate stiffness modulates regional stiffness distribution in iBMEC monoculture.**

Histograms of AFM-derived Young’s modulus values in IMR90-1 iBMECs cultured on substrates with stiffness of 1, 2.5, 15, and 194 kPa. Each panel represents relative frequency distributions measured at tricellular junctions (left), bicellular junctions (middle), or cell bodies (right). The final row shows overlaid distributions for all stiffness conditions.

**Figure S2. Substrate stiffness modulates regional stiffness distribution in iBAP co-culture.**

Histograms of AFM-derived Young’s modulus values in IMR90-1 iBMECs co-cultured with astrocytes and pericytes on substrates with stiffness of 1, 2.5, 15, and 194 kPa. Each panel represents relative frequency distributions measured at tricellular junctions (left), bicellular junctions (middle), or cell bodies (right). The final row shows overlaid distributions for all stiffness conditions.

**Figure S3. Substrate stiffness modulates regional stiffness distribution in iBAP without BC treatment.**

Histograms of AFM-derived Young’s modulus values in IMR90-1 iBMECs from iBAP group without any treatment. Each panel represents relative frequency distributions measured at tricellular junctions (left), bicellular junctions (middle), or cell bodies (right). The final row shows overlaid distributions for all stiffness conditions.

**Figure S4. Substrate stiffness modulates regional stiffness distribution in iBAP with BC treatment.**

Histograms of AFM-derived Young’s modulus values in IMR90-1 iBMECs in iBAP with MDA-MB-BR cell treatments. Each panel represents relative frequency distributions measured at tricellular junctions (left), bicellular junctions (middle), or cell bodies (right). The final row shows overlaid distributions for all stiffness conditions.

**Figure S5. Identification of stiffness-associated gene clusters in iBMEC monoculture and iBAP co-culture.**

Gaussian mixture modeling identified 36 stiffness-dependent gene expression patterns across four stiffness conditions in both monoculture (A) and co-culture datasets (B).

**Figure S6 Pattern-based clustering reveals stiffness-dependent gene expression trajectories in iBMECs and iBAP co-cultures.**

(A) Heatmaps showing Z-score–normalized expression of stiffness-responsive genes in IMR90-1 iBAP co-cultures across four substrate stiffnesses (1, 2.5, 15, and 194 kPa). Genes were grouped into distinct clusters (n = 2437, 1924, 441, 872, 926) based on shared expression trajectories using Gaussian mixture modeling. Each cluster displays a characteristic stiffness-dependent expression pattern (middle panels), while the right heatmap summarizes the top enriched Gene Ontology (GO) biological processes per pattern, with enrichment scores represented by color intensity. (B) Equivalent analysis for iBMEC monocultures, showing five dominant expression clusters (n = 2309, 512, 716, 523, 2079) with distinct stiffness-dependent trajectories.

**Figure S7. Representative genes related to stiffness-dependent transcriptional transition.**

Heatmap showing representative stiffness-responsive genes associated with cytoskeletal tension, mechanosensitive signaling, and junctional integrity.

**Table S1. Information of antibodies**

| Antibody | Vendor |  | Catalog number | Dilution |
| --- | --- | --- | --- | --- |
| ZO-1 | ThermoFisher |  | 61-7300 | 1:200 |
| GFAP | Abcam |  | ab68428 | 1:200 |
| PDGFR |  |  | Ab32570 | 1:200 |
| Goat anti-Mouse IgG (H+L) Cross-Adsorbed Secondary  Antibody,Alexa Fluor 488 | ThermoFisher |  | A11001 | 1:1000 |
| Goat anti-Mouse IgG (H+L) Cross-Adsorbed Secondary  Antibody,Alexa Fluor 568 | ThermoFisher |  | A11011 | 1:1000 |

**Video S1. Timelapse of cancer cell incorporation.**
