## Supplementary figures and images for "Substrate stiffness and cellular microenvironment regulate cell and junction mechanics in iPSC-derived brain microvascular endothelial cells"

### Supplemental figures

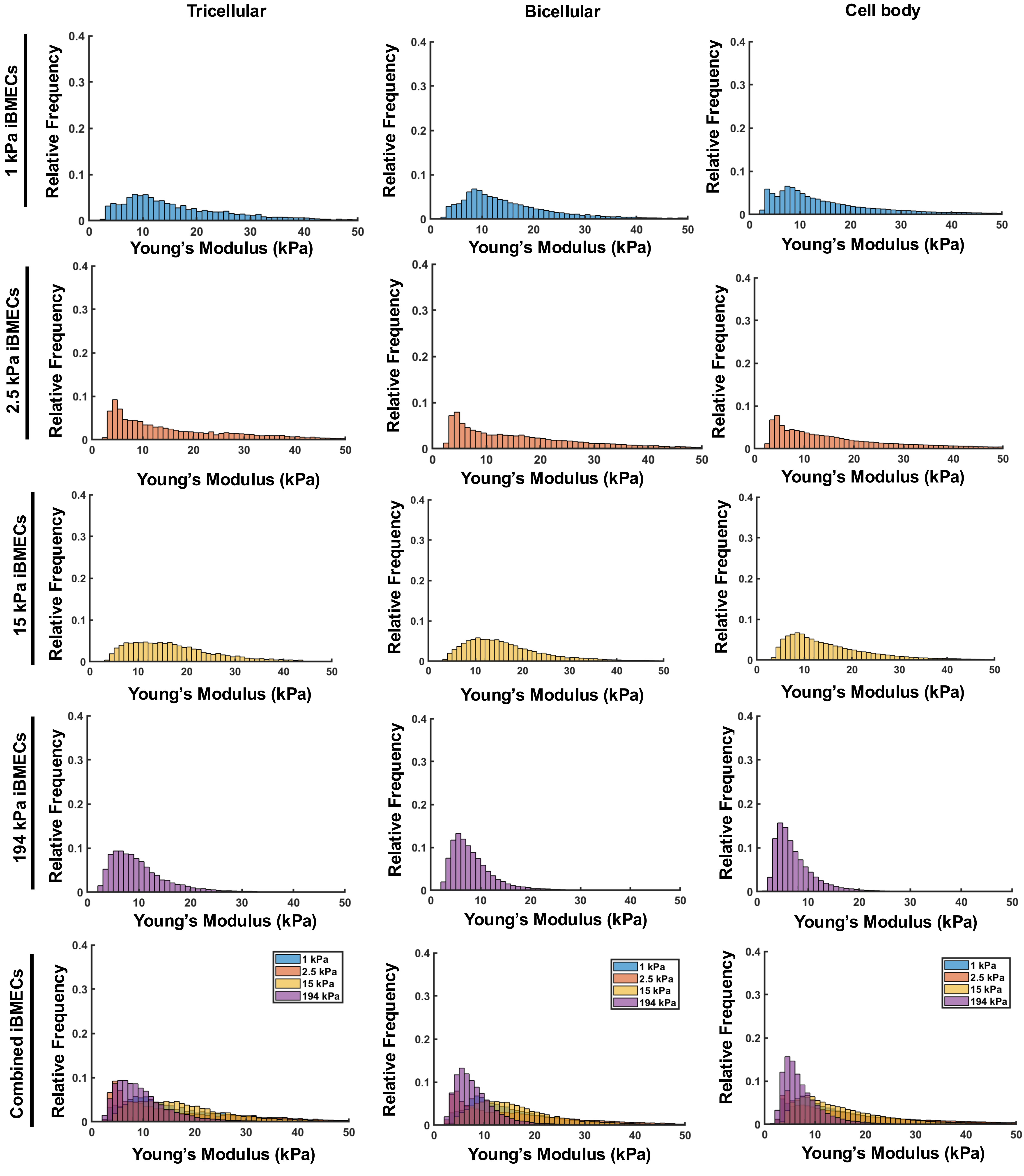

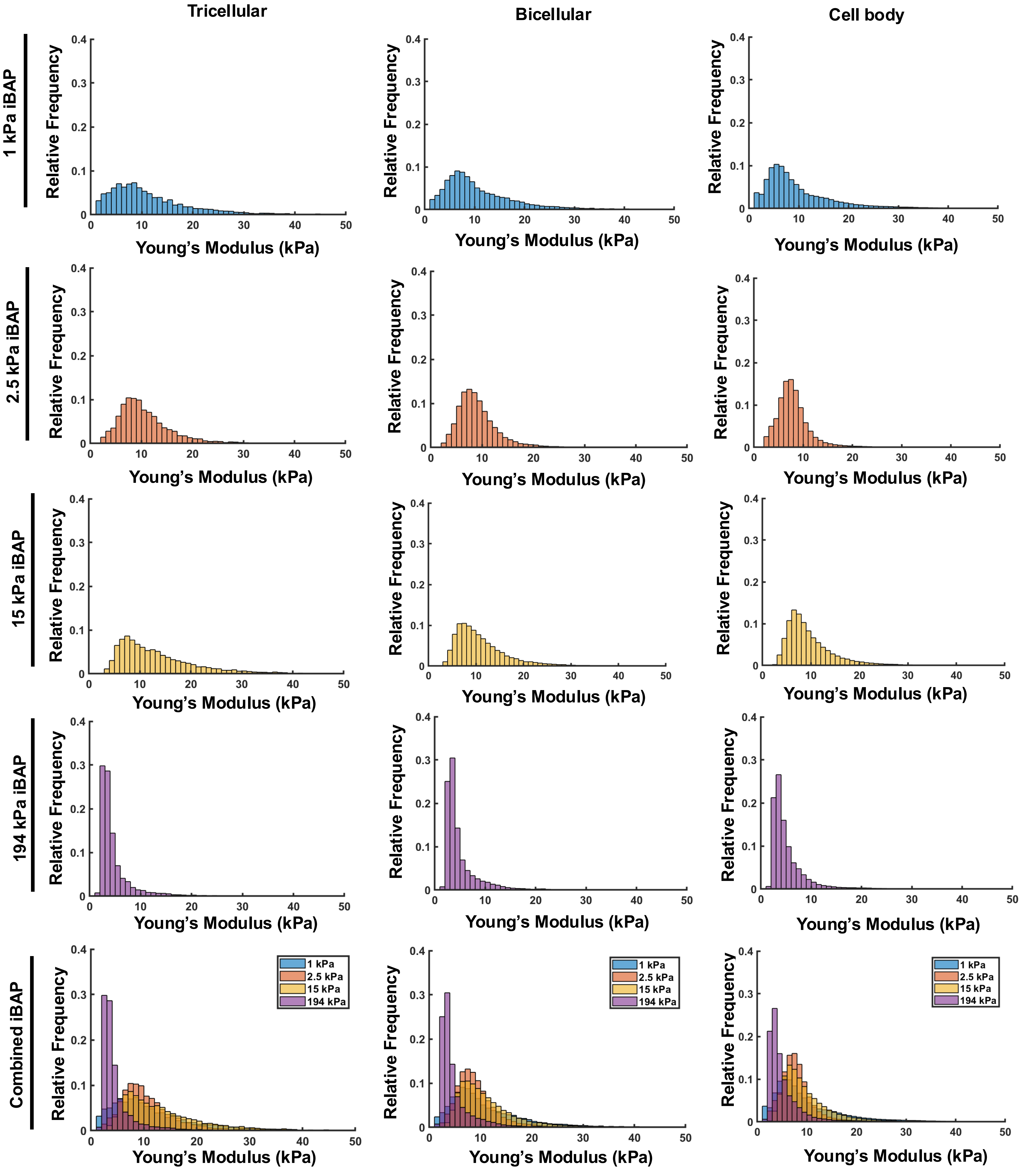

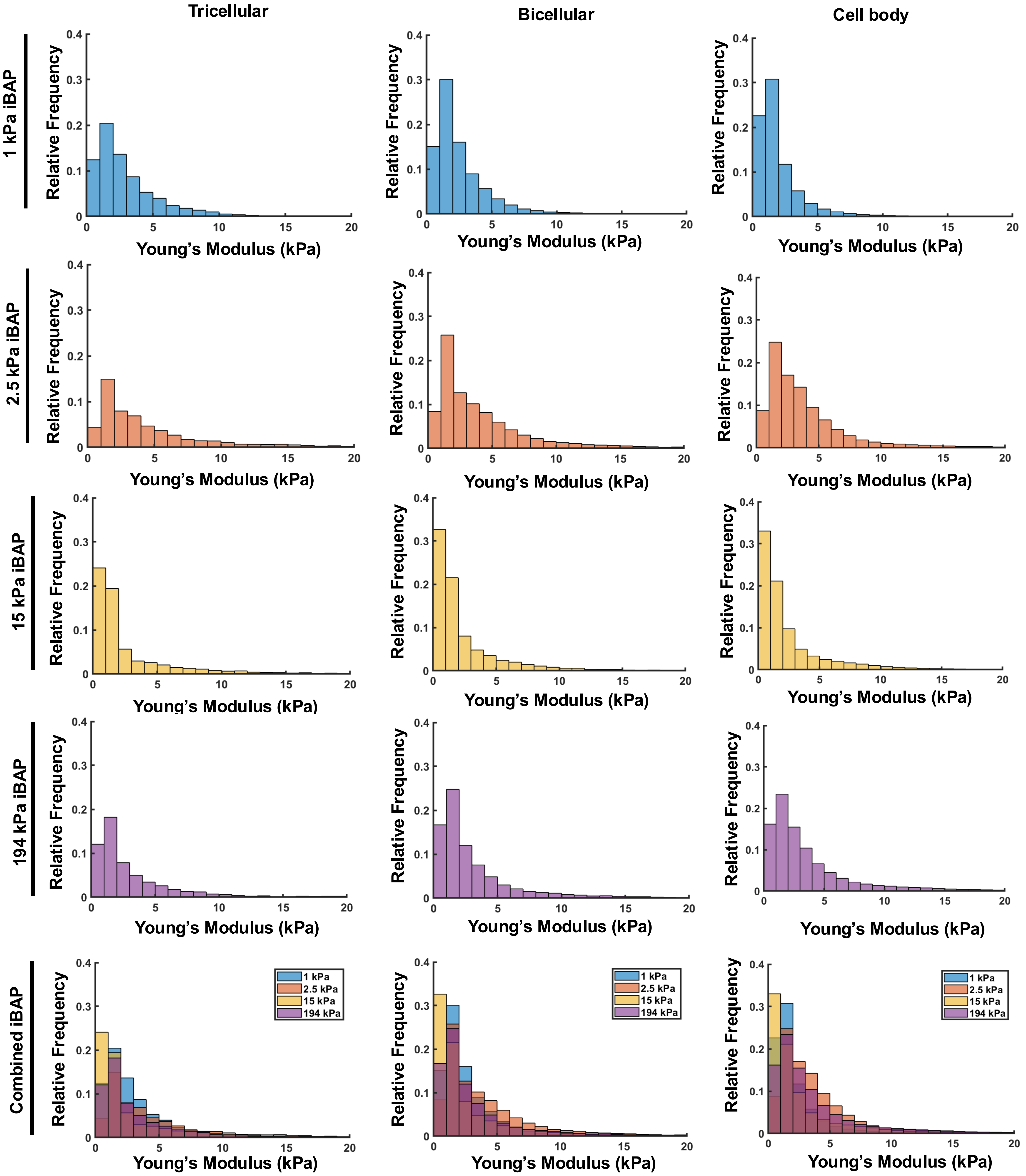

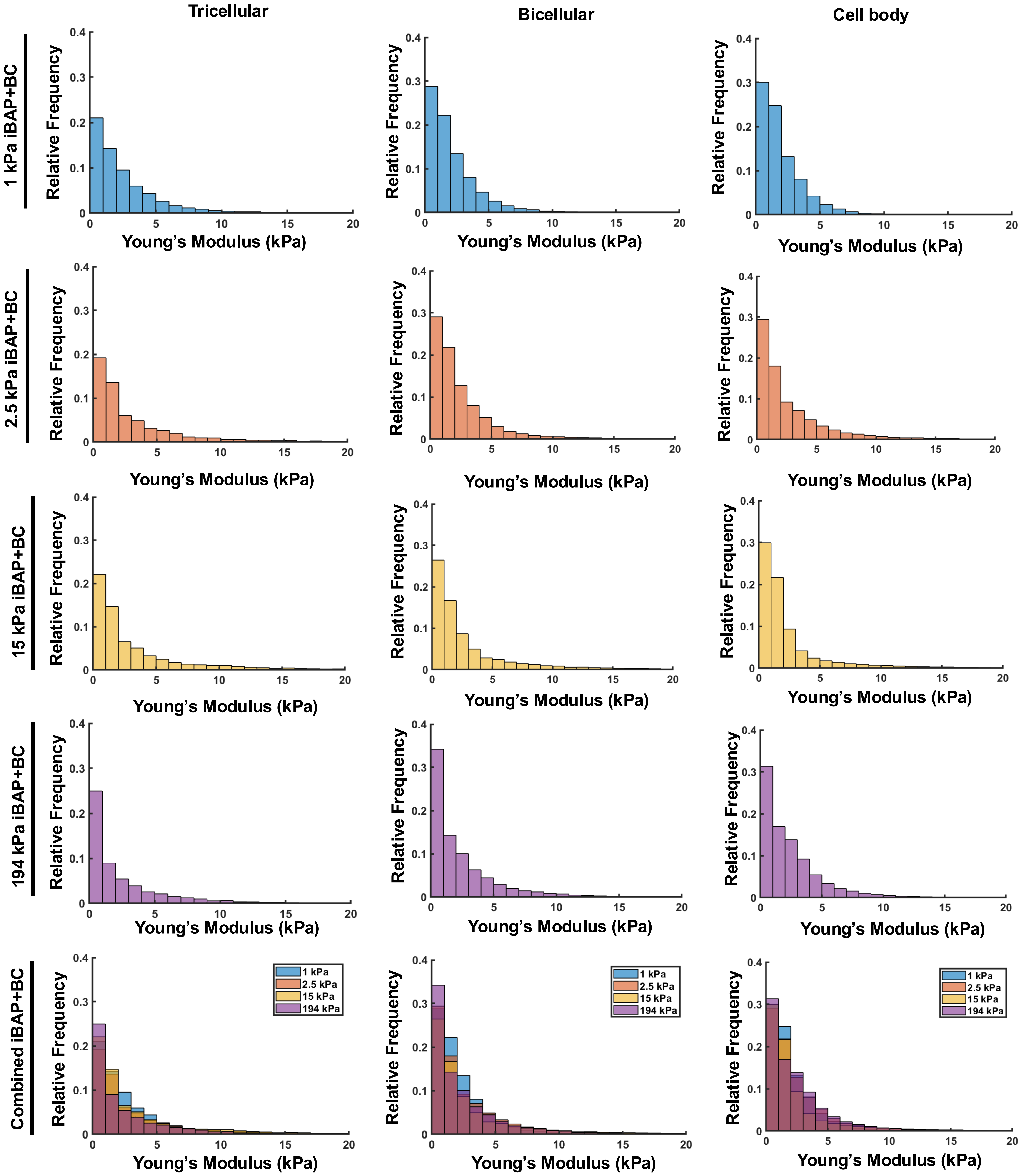

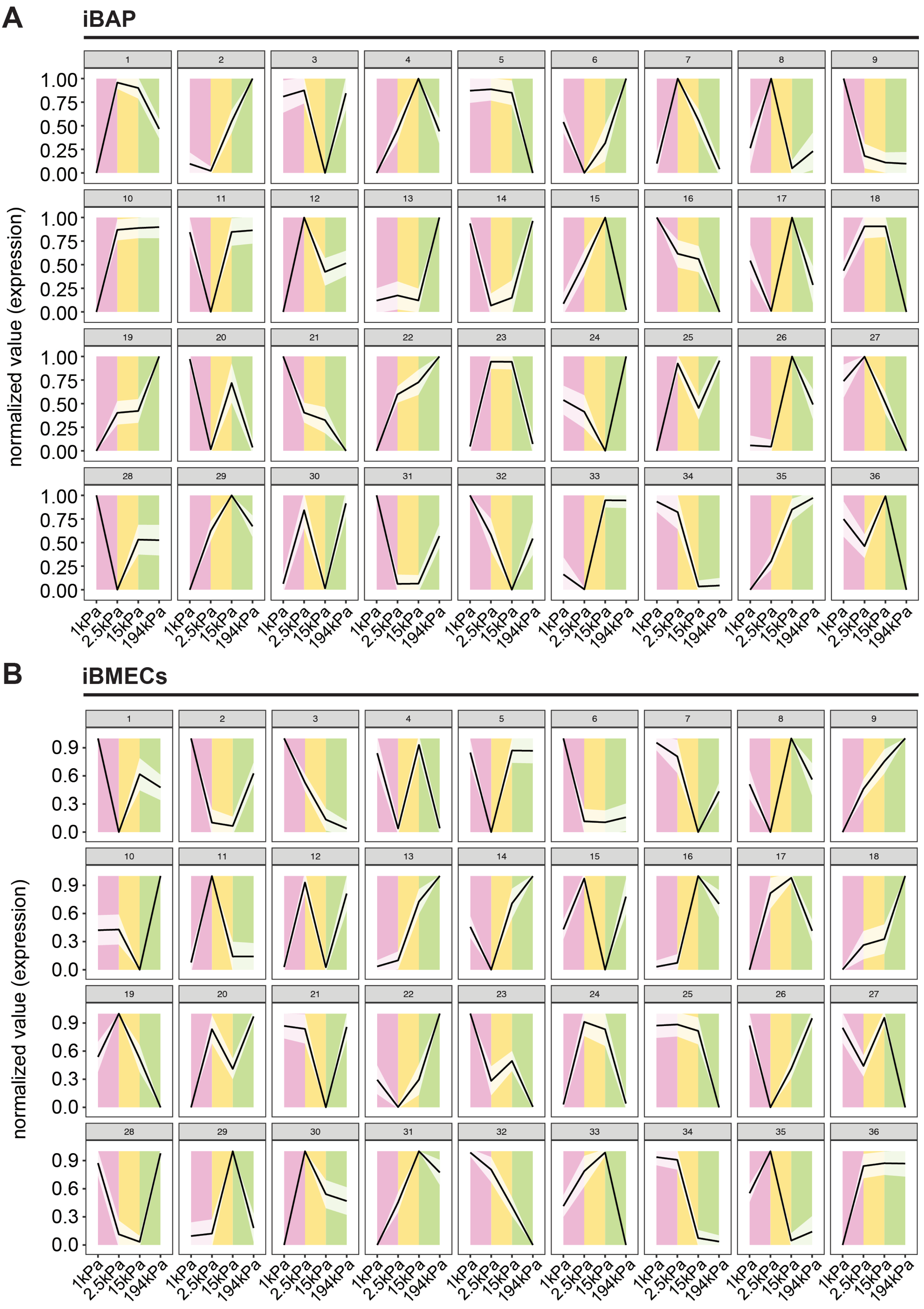

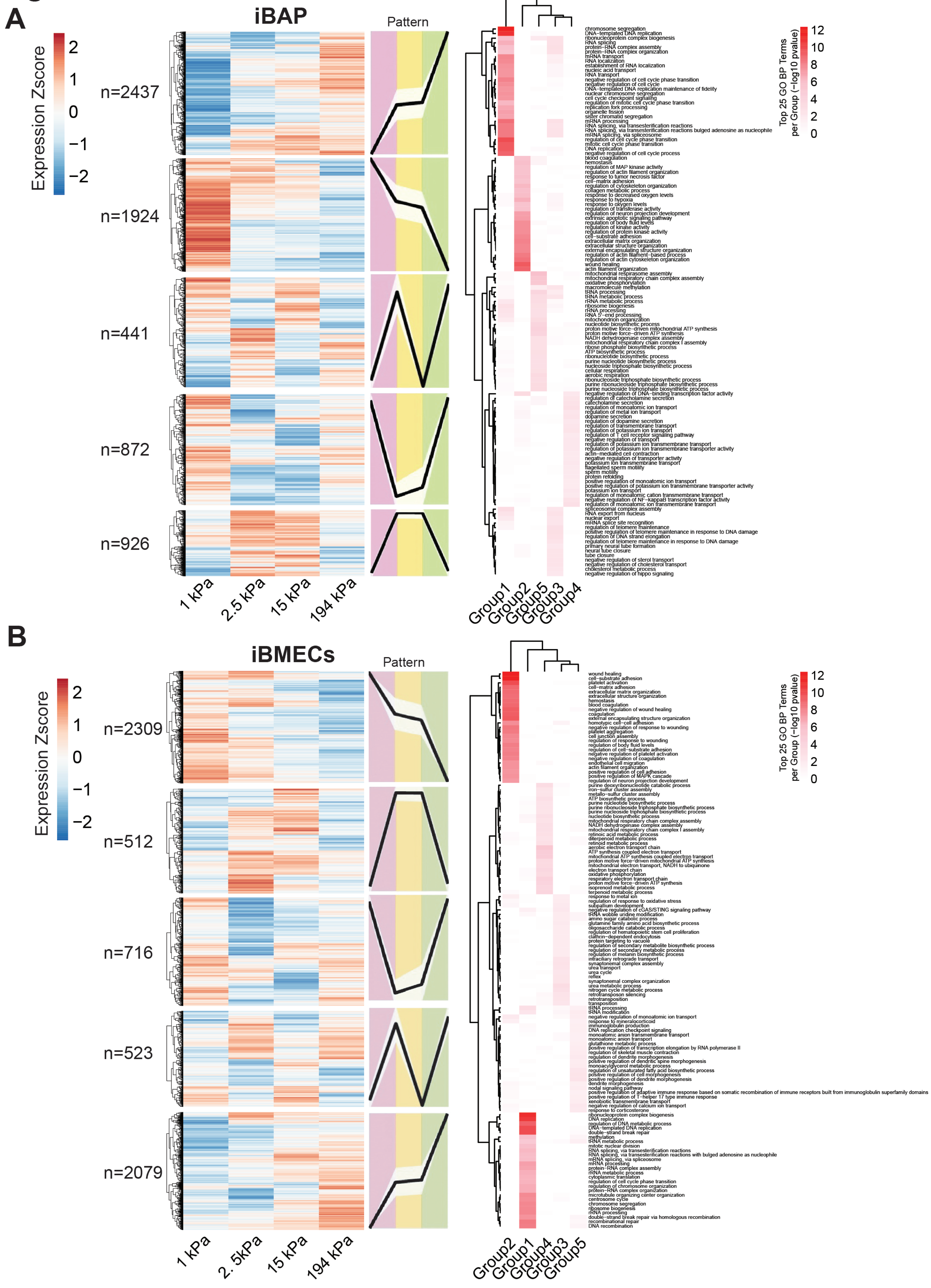

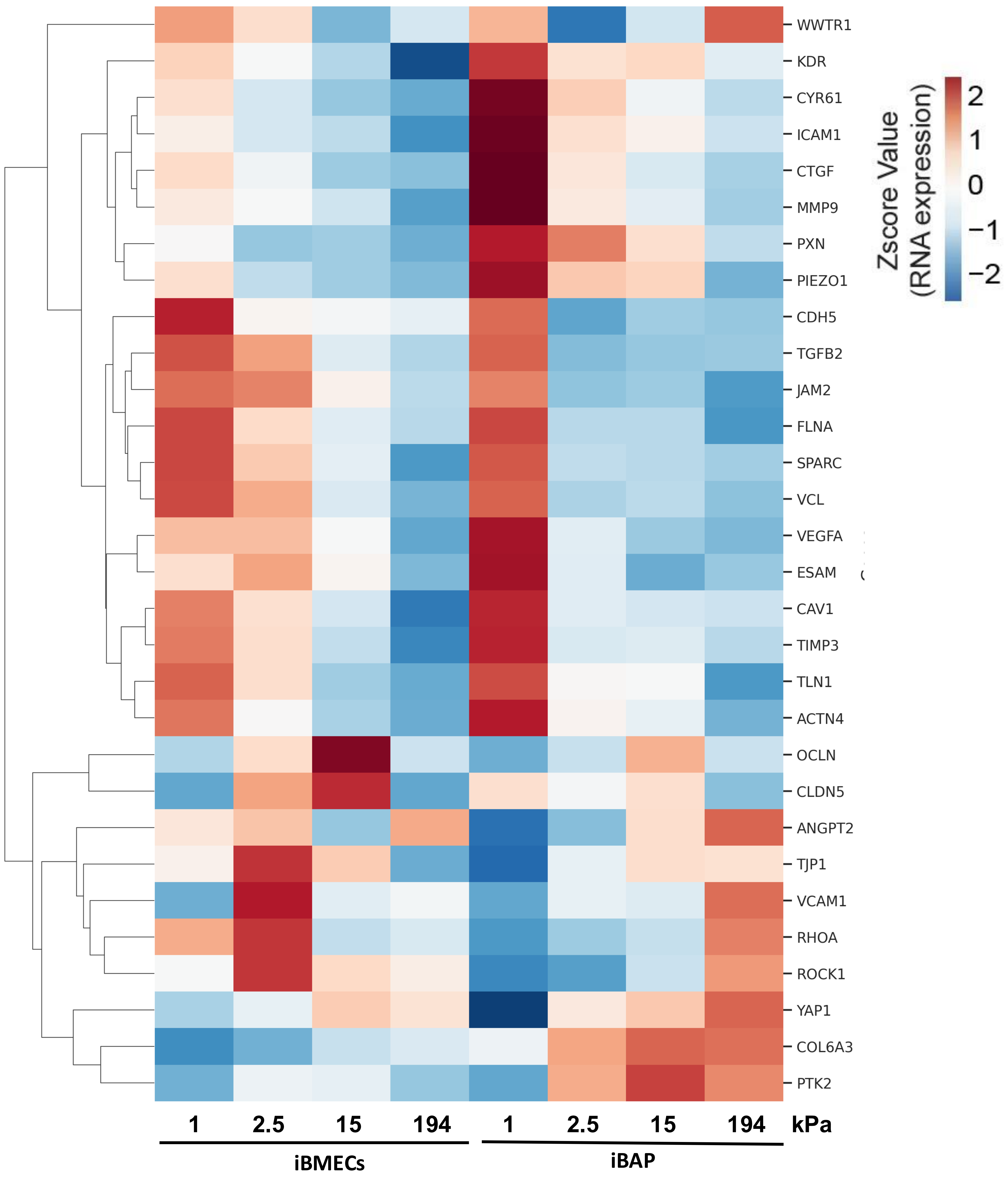
